# Modelling the invasion, management strategies and economic impact of the invasive alien aquatic plant *Egeria densa*

**DOI:** 10.1101/2024.08.29.610308

**Authors:** Adrián Flores-García, John Y. Dobson, Eva S. Fonfría, David García-García, Cesar Bordehore

## Abstract

*Egeria dens*a is a south American aquatic plant considered as an invasive alien species with a very aggressive and expansionist behaviour out of its native ecosystems. The widespread decorative use of *E. densa* in aquariums, coupled with the potential for subsequent disposal malpractice, has resulted in its introduction into a range of vulnerable ecosystems across the globe. This work presents a dynamic model of a population of *E. densa* based on its accidental introduction into an ecosystem in southeastern Spain. The model allows the simulation of colonisation process and management and disposal strategies. The model considers the influence of environmental variables such as temperature, light and water turbidity on the growth of the species. Among the management strategies tested, the optimal appears to be those of a preventive nature before the plant spreads, given that the costs of removal will be very high at that stage, and with high ecological impacts.

## 1. Introduction

Invasive alien aquatic plants (IAAPs), generally called exotic plants, are responsible for the destabilisation of ecosystems they colonise where they are not native. They are characterised by an often rapid and aggressive expansive dynamics as they normally do not encounter any other organism to moderate their development (Powell, et al., 2011; Reid et al., 2019). This is especially true in aquatic environments (Hussner, 2012; Vilà et al., 2010), where the spread of propagules and plant fragments are more difficult to control, having a greater probability of survival (Essl et al., 2018).

One of these IAAPs is the freshwater plant *Egeria densa (Planchon., 1849)* (Figure 1). Also known as brazilian elodea, brazilian waterweed and leafy elodea (Smith et al., 2022), it’s a submerged aquatic macrophyte species from the Hydrocharitaceae family native to South America, in particular from Minas Gerais (Brazil) to the River Plate delta in Argentina (Yarrow et al., 2009; Talavera & Gallego, 2010). It’s included in the Spanish Catalog of Invasive Exotic Species since 2013 (Real Decreto 630/2013), and also has been regulated at regional level in the Valencian Community to approve control measures for exotic species (Decreto 213/2009).

**Figure 1.**
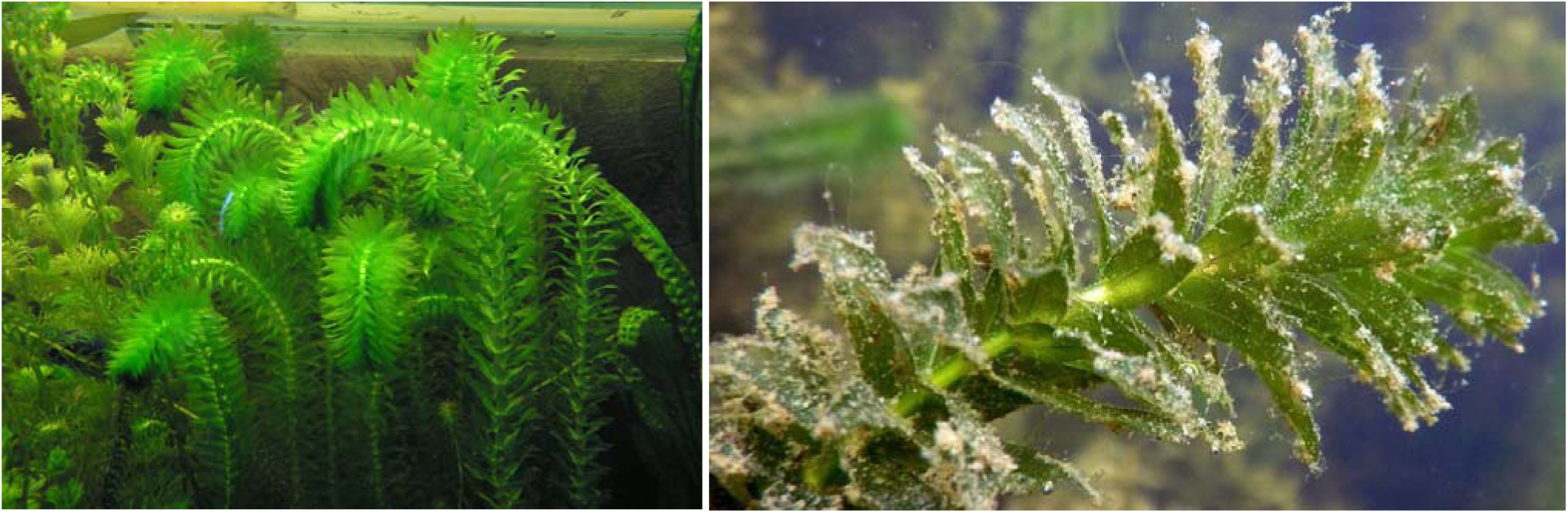
Ecosystem with large amounts of *Egeria densa (Planchon, 1849)* and the plant in detail. Source: Igor Sheremetyev - Plants of the World (https://powo.science.kew.org/taxon/urn:lsid:ipni.org:names:431710-1) and Wikipedia (https://es.m.wikipedia.org/wiki/Archivo:Egeria_densa_%C3%A9tang_de_Saint-Nolf_14_ao%C3%BBt_2015a7.JPG).

The ecology includes perennial nature, living submerged (Getsinger & Dillion, 1984), surviving the winter in the form of shoots, and the capacity to change size over the course of the year as an adaptation to store starch during winters and shrinking when not needed (Figure 2) (Haramoto & Ikusina, 1988; Yarrow et al., 2009). It is a dioecious species that reproduces sexually when conditions are favourable. The reproduction can be either sexual, when the pollen reaches the female eggs (Getsinger & Dillion, 1984), or asexual by fragmentation, when a fragment takes root in the substrate, giving rise to a new individual (Bonilla-Barbosa & Santamaria, 2013). The aggressive invasive behaviour of *E. densa* is based on a marked adaptation to a variety of ecosystems, in particular the ability to grow in low light and cold temperatures (Getsinger & Dillion, 1984; Curt et al., 2010), as well as the flexibility to absorb nutrients from the water column and sediments (Conrad et al., 2016; Durand et al., 2016). It is considered by several authors as a plant ecosystem engineer, due to the way it changes the physical properties of the environment: temperature, raising oxygen (Smith et al., 2022), nutrients, hydrodynamics or phytoplankton reduction, among others (Jones et al., 1994; Jones et al., 1997; Yarrow et al., 2009). It is widely used as a decorative aquarium plant all over the world, causing its expansion after the release into the environment (Yarrow et al., 2009; Lansdown, 2011; Osaki and Kodama, 2017), a possible reason for its presence in the waters of virtually every continent (CABI, 2020).

**Figure 2.**
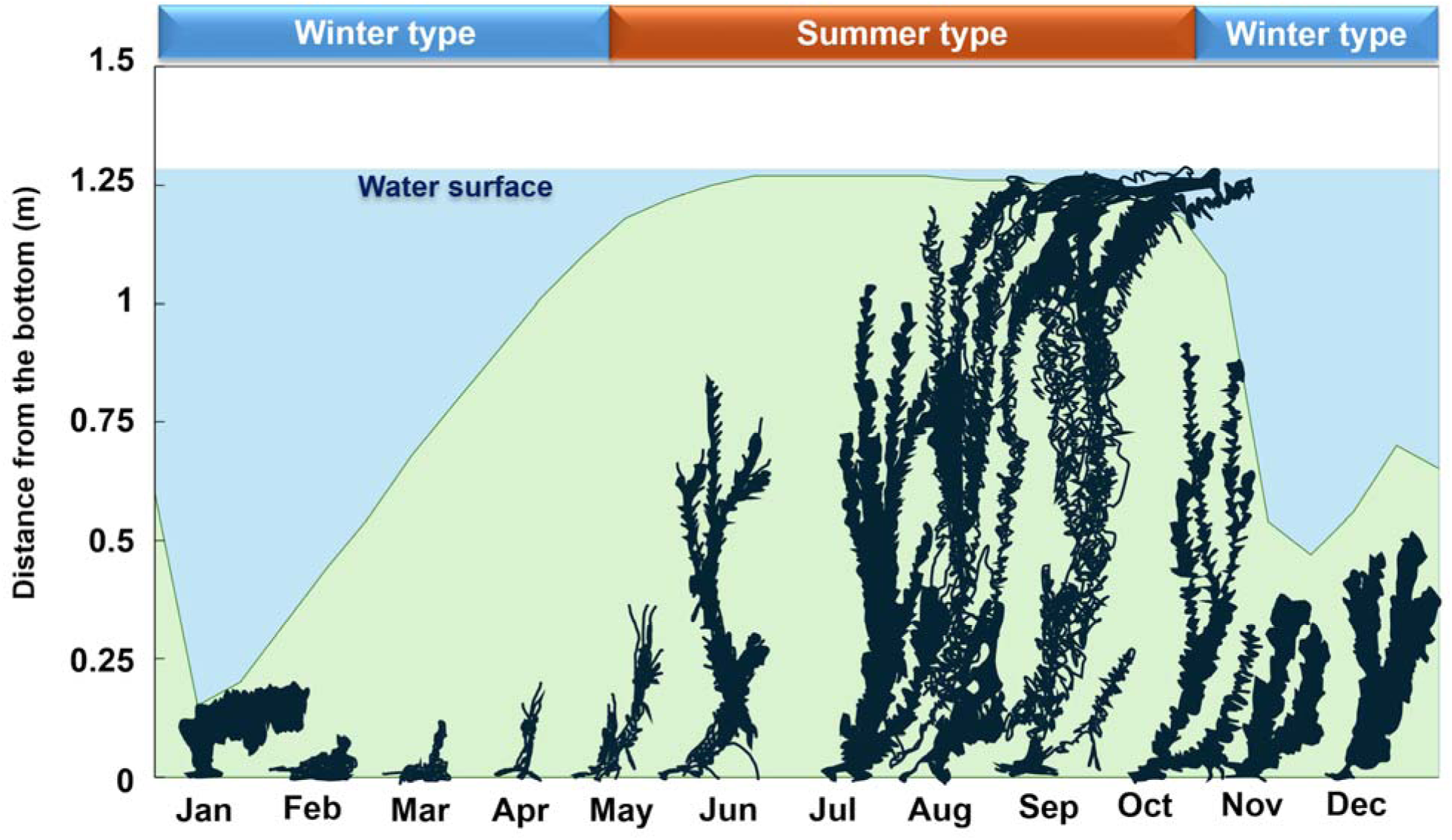
Seasonal changes in *Egeria densa* size. Adapted from Haramoto & Ikusina, 1988.

In Spain (Figure 3), *E. densa* was first identified in the early 1900s in the Retiro Park (Madrid). It was determined that the specimen was misidentified as another species, although other authors have confirmed that it has been collected since 1912 (Medina & Bracamonte, 1995; Talave a & Gallego, 2010). Subsequently, the species was detected in the 1990s in Gandia, in the Valencian region (Cirujano & Medina, 1995) and in various locations in Catalonia (Gutiérrez & Sáez, 1996; Bolòs & Vigo, 2001). Later in the 2000s was found in continental waters in Galicia (Pulgar & Izco, 2005), reservoirs and lakes in the Basque Country (Aizpuru et al., 2000), marshes in the province of Seville (Gros et al., 2009), and the Pego-Oliva marsh (South of Valencia province and North of Alicante province) according to several media reports. More recently, it has been detected in new locations in Catalonia (Aymerich, 2012) and La Palma (Canary Islands) (Otto & Verloove, 2020).

**Figure 3.**
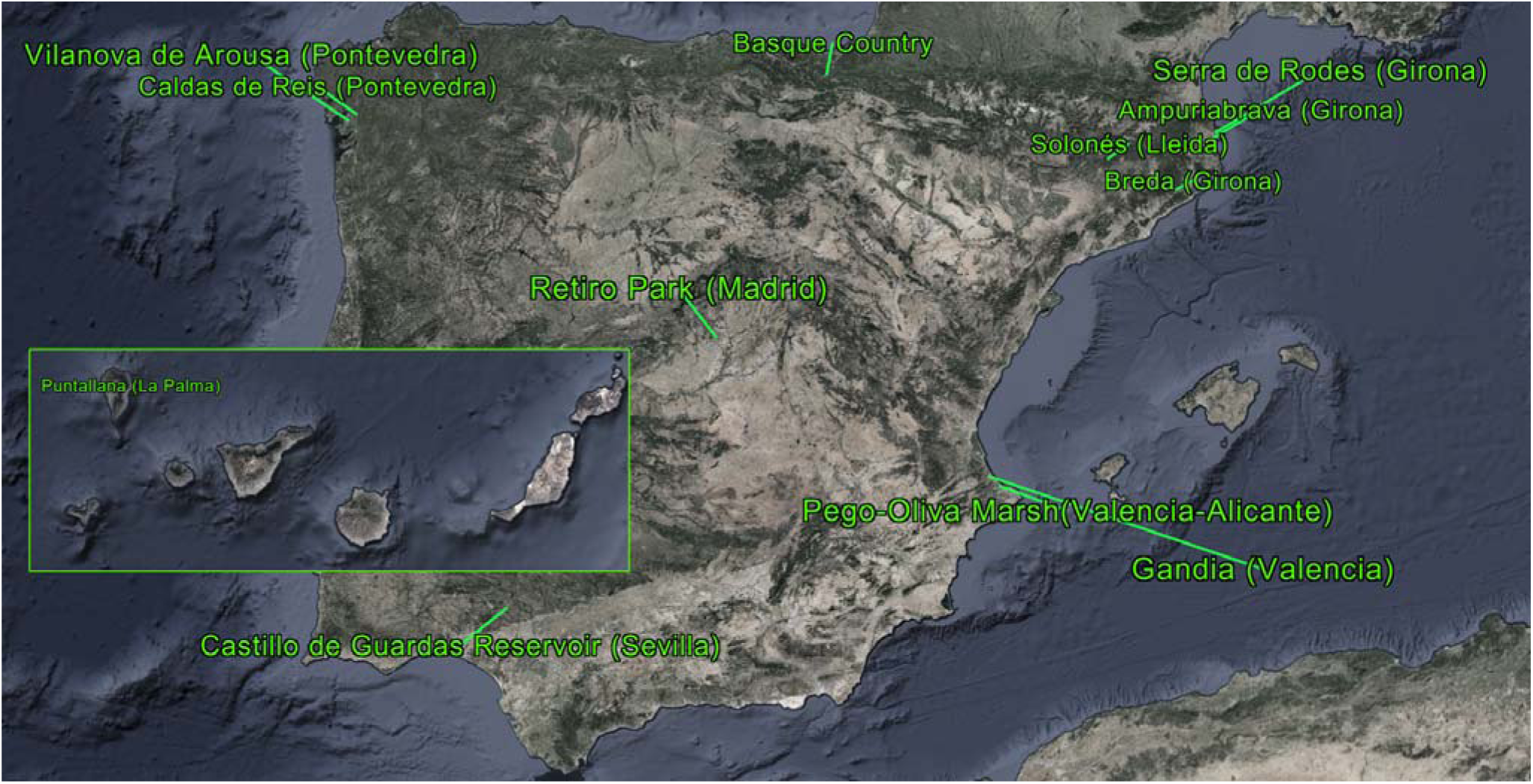
Locations in Spain where *Egeria densa* has been observed since the 1900s. Source: own elaboration.

*E. densa* impacts are described as a drastic alteration of the ecosystem characteristics: prevents the entry of light affecting other species, increases oxygen levels, sequesters nutrients, reduces habitat size for other species, slows down water courses, and alters the trophic networks, leading to competition for food and other negative effects on native species (Yarrow et al., 2009; Darrin et al., 2009; CONABIO, 2014). *E. densa*, as other invasive plants, can displace other species to less favourable environments or even make them disappear in an indirect way (Wells & Clayton, 1991). In terms of economic and social impact, the plant has the potential to cause significant inconvenience to fishing, swimming, boating and other aquatic activities. High densities of *E. densa* have been observed to even cause blockages in hydroelectric plants in Brazil and New Zealand (Barreto et al., 2000; Darrin et al., 2009). In relation to human health, it does pose an indirect danger in terms of the spread of *Anopheles* mosquitoes (Johnson et al., 2006; Pennington & Sysma, 2009) and even the death by drowning due to being trapped in high densities (García, 2019). Wide range of methods against the expansion of the species are documented, but no effective methods have been identified for its total eradication. Considering this, most plans and efforts are focused on the prevention of establishment and minimise the spreading (Meacham, 2001; Johnson et al., 2006).

This research’s aim consists of the development of a dynamic model to recreate the life cycle of *Egeria densa* and the colonisation process, along with environmental variables. In addition, the model serves as a way to test different management strategies and their cost, considering the environmental effects and crop reduction.

## 2. Data

### 2.1. Study site

We chose the Pego-Oliva marsh area (38° 52’ 16’’N, 0° 3’ 39’’W) as an example to model a colonised ecosystem. The total area of this marsh is 1,290 ha, of which we will assume 10% is water, the rest being mainly agricultural fields for orange trees. The model will consider the water canals with an area of 129 hectares, a mean of 1.5 metres of water depth, and approximately a third part dedicated to rice crops, according to official data where 360 ha were established as crops (Generalitat Valenciana, 2021).

### 2.2. Study species

An exhaustive bibliographic search about the biology and ecology of *E. densa* was conducted in specialized databases such as Google Scholar, Sciencedirect and Dialnet to feed the model with the values of the needed parameters (Table 1). Some factors that positively influence the growth of *E. densa* are higher temperatures, low water turbidity, low salinity, high O_2_ concentration and low nitrogen and phosphorus levels. Special importance is given to temperature and turbidity (Lacet et al., 2019).

**Table 1.**
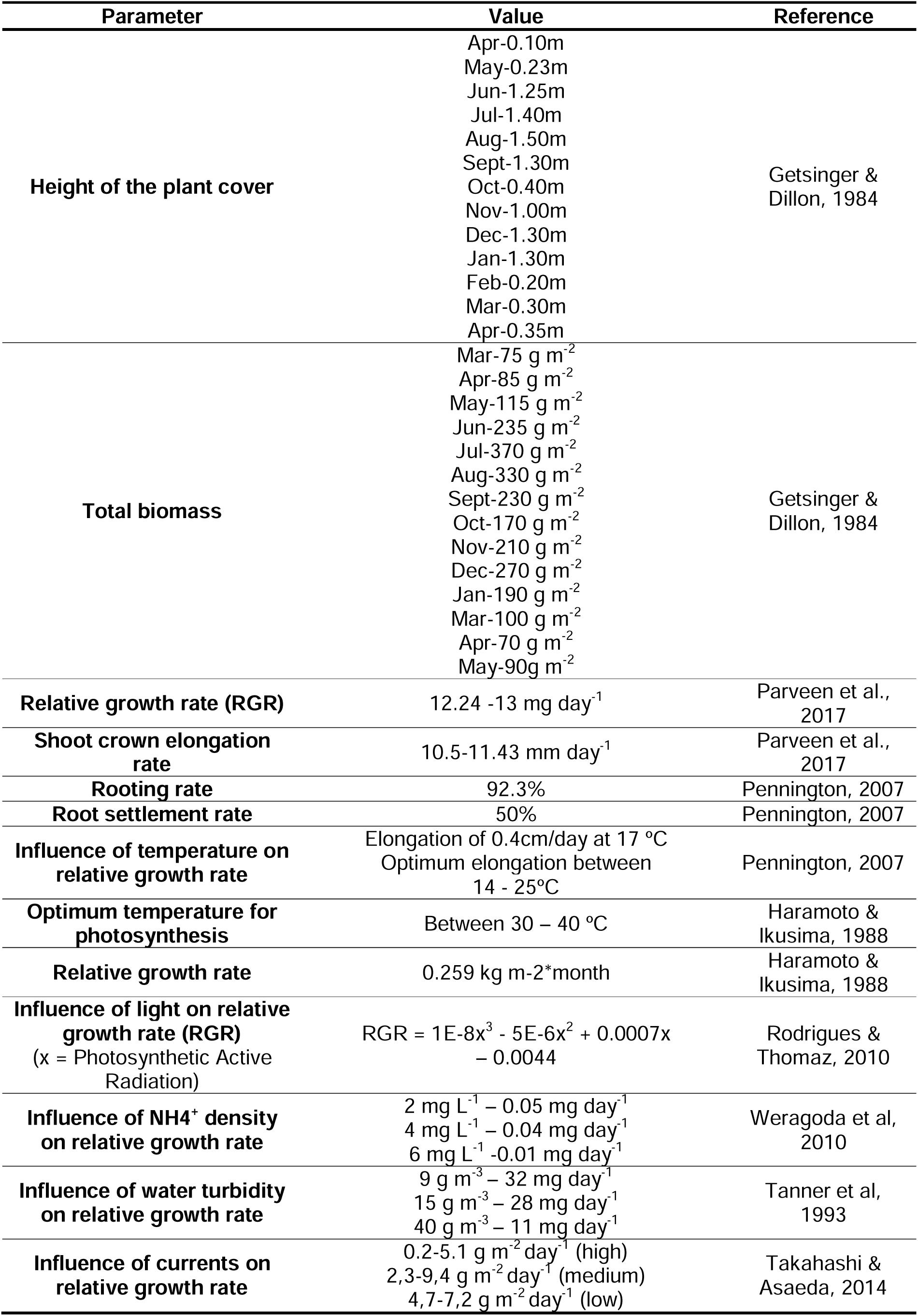
*Egeria densa* parameters applied in the dynamic model that were obtained from previous research.

## 3. Dynamic model

### 3.1. Model structure

We used the software STELLA Architect (Isee Systems) to build the *E. densa* model. This program is based on differential equations, and it’s shown as a friendly and intuitive environment composed of stocks, connectors, and converters. Stocks are the building blocks that store, in this case, *E. densa* biomass. These stocks are linked thanks to connectors, which enable them to fill, drain or move accumulations between them. Then, converters introduce constants, external inputs, equations, or graphical functions which control the flows and define the dynamics of the model.

The main part of the model is composed of three stocks: crown shoots, planted *Egeria* and floating *Egeria* (Figure 4). The *crown shoots* stock represents every shoot that will become a functional stem in the following stock. It has three input flows, the first of which comes from species’ sexual reproduction, depending on the *number of flowers* and a *fecundation rate*, and the other two come from asexual crown shoots generated asexually by planted or floating *Egeria*. The flow connecting with the next stock is modulated by a *shoot growth rate* and a *seasonal pattern*.

**Figure 4.**
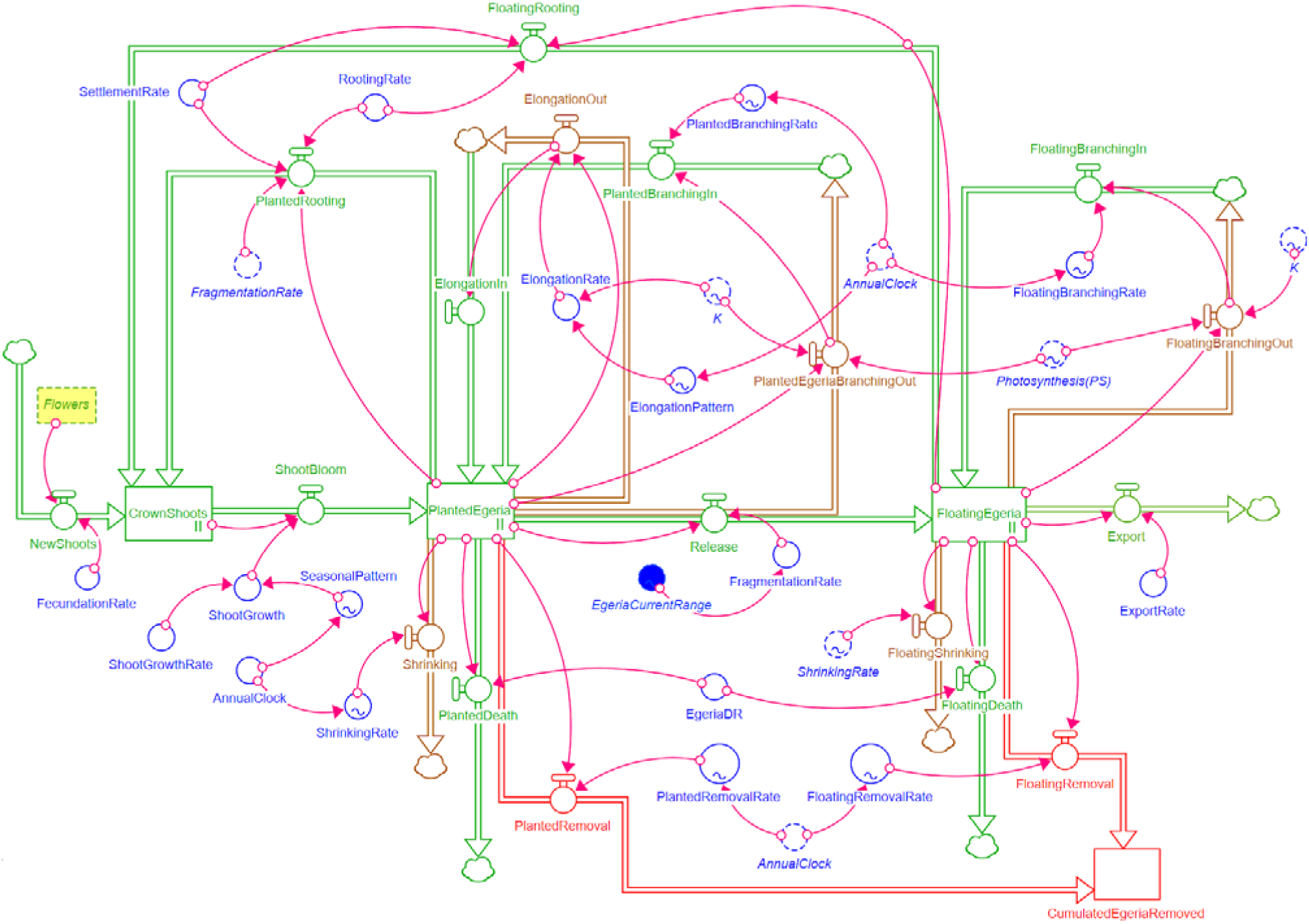
*Egeria densa* model main module which represents the plant’s life cycle.

Then, the *planted Egeria* stocks represent the plants that remain attached to the bottom. Here we consider the plants’ biomass growing, as elongation and branching, and the loss in the form of seasonal shrinking and death. Elongation and branching depend on two parameters which change every month of the year, these are *Elongation Rate (ER)* (coming along with an *Elongation Seasonal Pattern*) and *Planted Branching rate (PBR).* Death exit flow is determined by an *Egeria Death Rate (DR)*. Shrinking is the seasonal variation of plant’s size, being bigger when the temperatures are lower and smaller during higher summer temperatures. This exit flow is modulated by a *shrinking rate (SR)* that changes its value depending on the month. The last exit flow continues into the last stock and represents the fragmentation of the apical ends of touching-bottom *E. densa* due to environmental water currents.

*Floating Egeria* stock represents the plants’ fraction that is no longer attached to the bottom. The entry into this stock is managed by a *Fragmentation Rate (FR)* which controls the release of *E. densa* planted branches. Moreover, in this stock we still consider plants’ shrinking and death, both modulated by the same parameters as planted *Egeria*, except for branching, which has its own *Floating Branching Rate (FBR),* and the addition of the possibility of *E. densa* exit of the system as an export, modulated by and *Export Rate (ExR)*.

As a dioecious species, it has two different reproduction mechanisms (sexual and asexual). Both are represented in the model (Figure 4 as rooting rates and flows, and Figure 5 by sexual reproduction). The sexual mechanism consists of the reproduction by flowers, where the pollen fertilises the female eggs, and this results in a propagule which will end up transforming it into new shoots.

**Figure 5.**
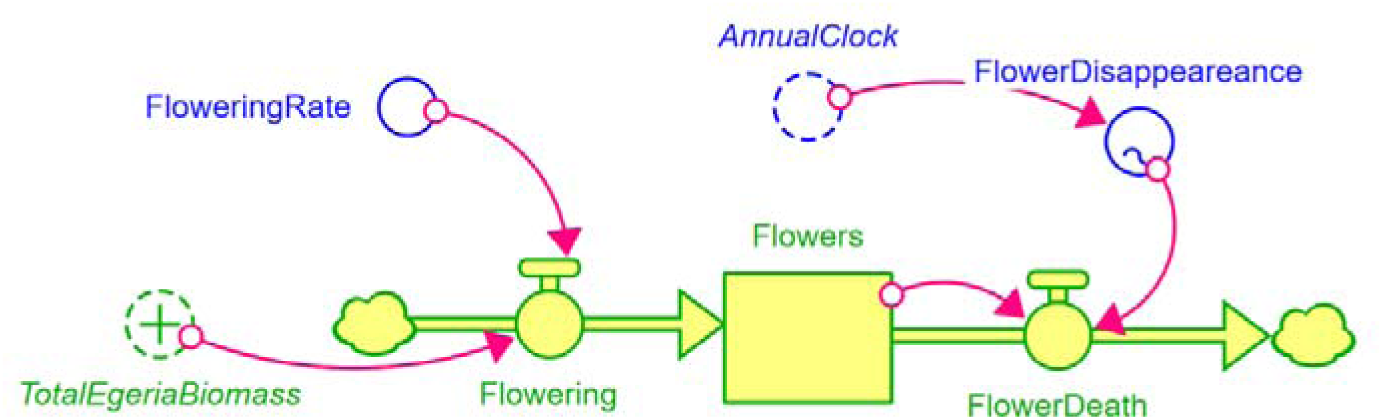
*Egeria densa* flower stock and the variables that represent the sexual reproduction of the plant.

The asexual reproduction is the strategy that *E. densa* uses the most e.g. fragmentation. Part of the stem releases roots that may reach the bottom and establish. In the model, this can happen to both planted and floating plants stocks, regulated by a *Rooting Rate (RR)* that controls two flows whose destination is the crown shoot stock. This happens for both planted and floating stocks (see Figure 4 *PlantedRooting* and *FloatingRooting* flows).

Other characteristics of the model are:

1. Additional flows and variables dedicated to exploring the options for the removal of *E. densa* from the environment (see red flows from the underside of Figure 4). This allows us to remove the amount of either *planted* or *floating E. densa* whenever we define the months of the year.
2. *E. densa* carrying capacity sub-model (Figure 6). Even with the perfect conditions for the plants’ growth, the population can’t grow to infinity. We first use a *density dependent carrying capacity (K)* that acts as a limit of plant’s growth. This *K* is calculated by the formula (see Eq. 1):

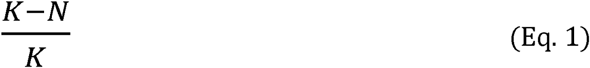

Biomass per cubic meter is calculated by dividing the sum of biomass of *Planted Egeria* and *Floating Egeria* stocks by the 129 water hectares and dividing again by the 1.5 meter *water depth* of the study area. This *K* limitation is being applied in the plant’s *elongation* and *branching* parameters, and we set up a maximum density of *E. densa* around 2 kg m^-3^, according to field data.
3. A set of environmental variables affect *E. densa* life cycle (Figure 7). This includes a variable input connected to a graphical parameter that defines how the variable influences the plant. Environmental physical and chemical variables considered are light, temperature, ammonia, turbidity and water currents. The values and how they influence plants’ growth are taken from previous research (see Table 2). All these variables have an influence on photosynthesis, except for the water currents that will modulate the *Fragmentation Rate* (*FR*).
4. An economic module was set to consider the costs of the activities around the ecosystem management, such as species conservation, tourism or rice crops, and the measures against the expansion of *Egeria* (Figure 8). Contains numeric indicators of key variables which would facilitate an easy read of the outputs.

**Figure 6.**
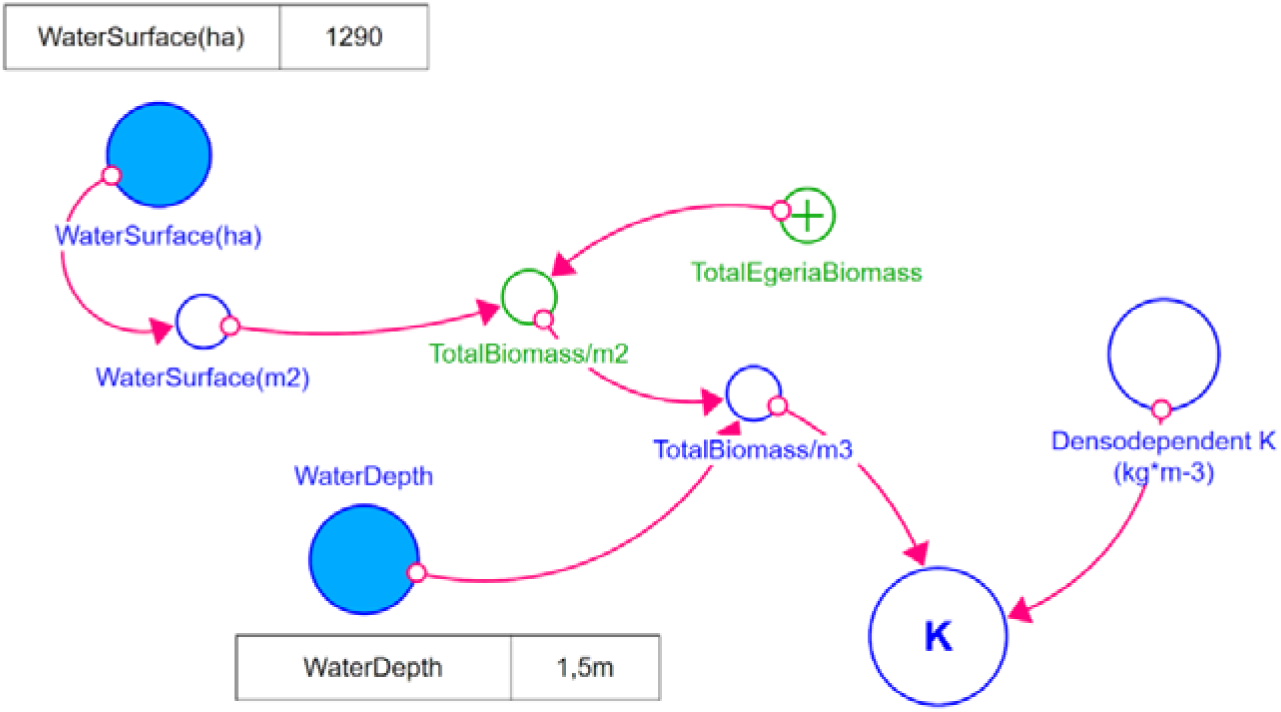
*Egeria densa* carrying capacity set of variables.

**Figure 7.**
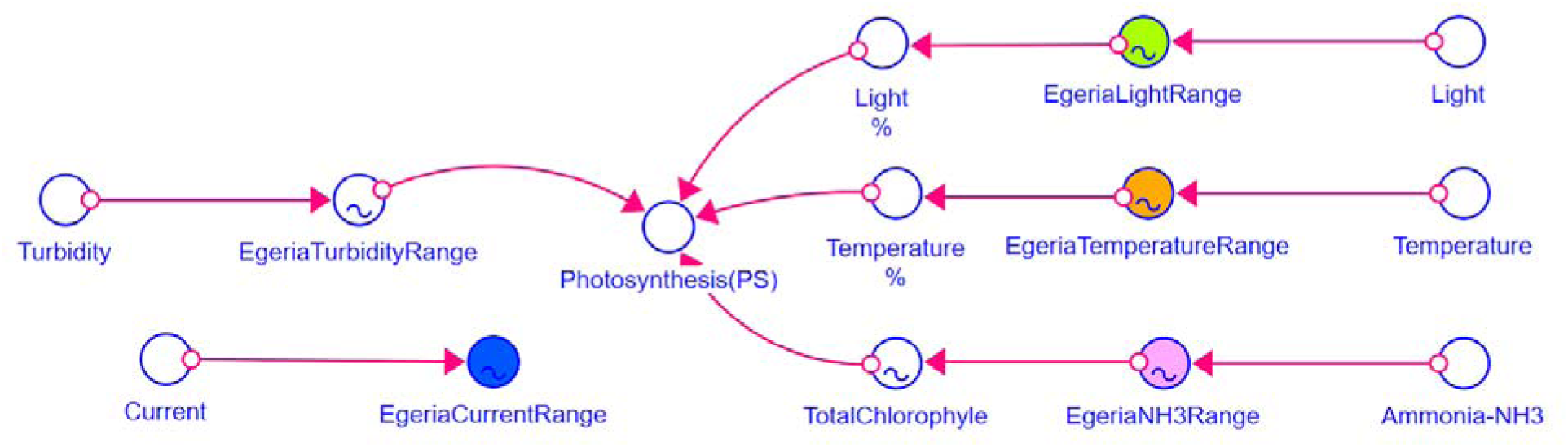
*Egeria densa* influencing environmental variables and their inputs.

**Figure 8.**
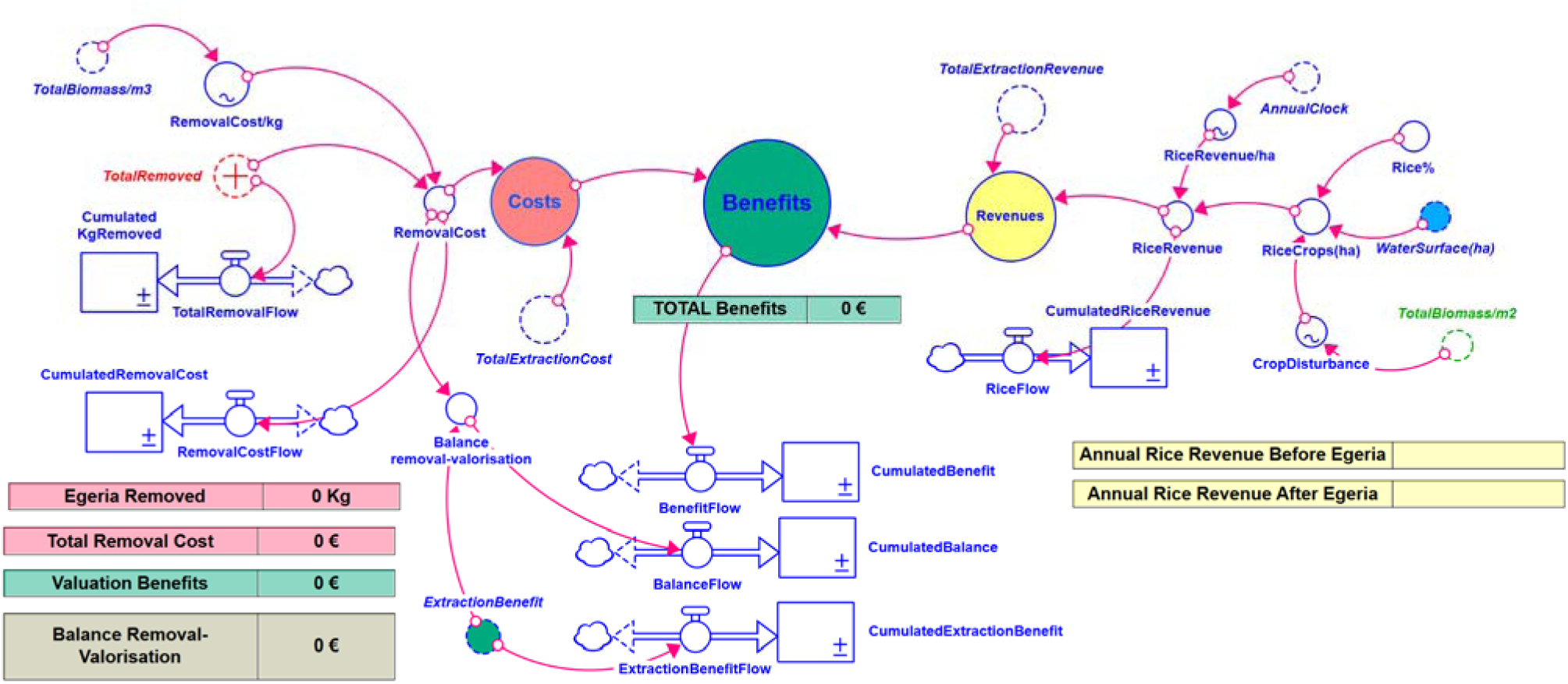
Economic module of the *E. densa* model with every activity involved in the economy of the system.

The main output is a benefit variable where the value of the ecosystem can be estimated along the possible impact that *E. densa* has. The Stella Architect file and the script of the model can be downloaded from Supplementary Material at https://bit.ly/4cIzZOa.

Almost half of this economic module collects a very simplified version of rice’s agricultural activity in the study area, showing revenues, costs and benefits, seasonality and disturbance associated with *E. densa*. This part of the model will allow us to explore the effect of the invasive plant on agricultural activity revenues.

### 3.2. Model output

The base scenario simulates that a starting amount of 2 kilograms of *Floating Egeria* enters the 129 ha system (Figure 9). This emulates the release of an exotic species. The modelisation shows an exponential increase on *E. densa* biomass with higher increasing velocity between months 100 and 150, that is from year 8 to 12. This exponential growth rises plant’s biomass and density faster the following 11-12 years until the carrying capacity of the system is reached and makes the *E. densa* population fluctuate around the established 2 kg m^-2^ as long as continues the simulation.

**Figure 9.**
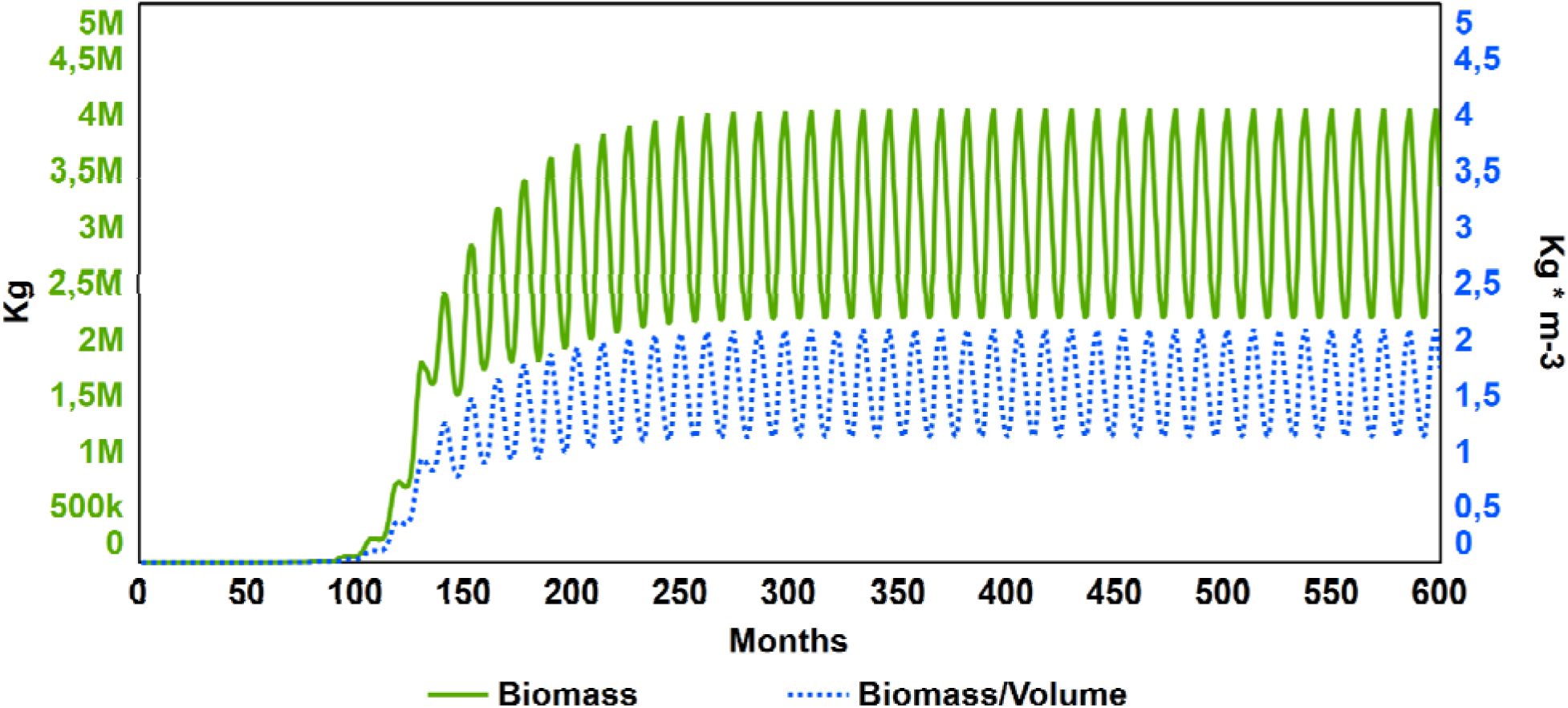
*E. densa* total biomass evolution after the introduction of 3 kg at time 1 during 50 years in the 129 hectare theoretical system.

In relation of the economic output (Figure 10), derived from rice crop benefits where we estimated a benefit per hectare of 1,750€, the profits decreased rapidly from year 10 to 12 from 74,500 to 21,200 € per year, reaching a steady state, when *E. densa* had extended all over the ecosystem reaching its maximum abundance (carrying capacity of the system).

**Figure 10.**
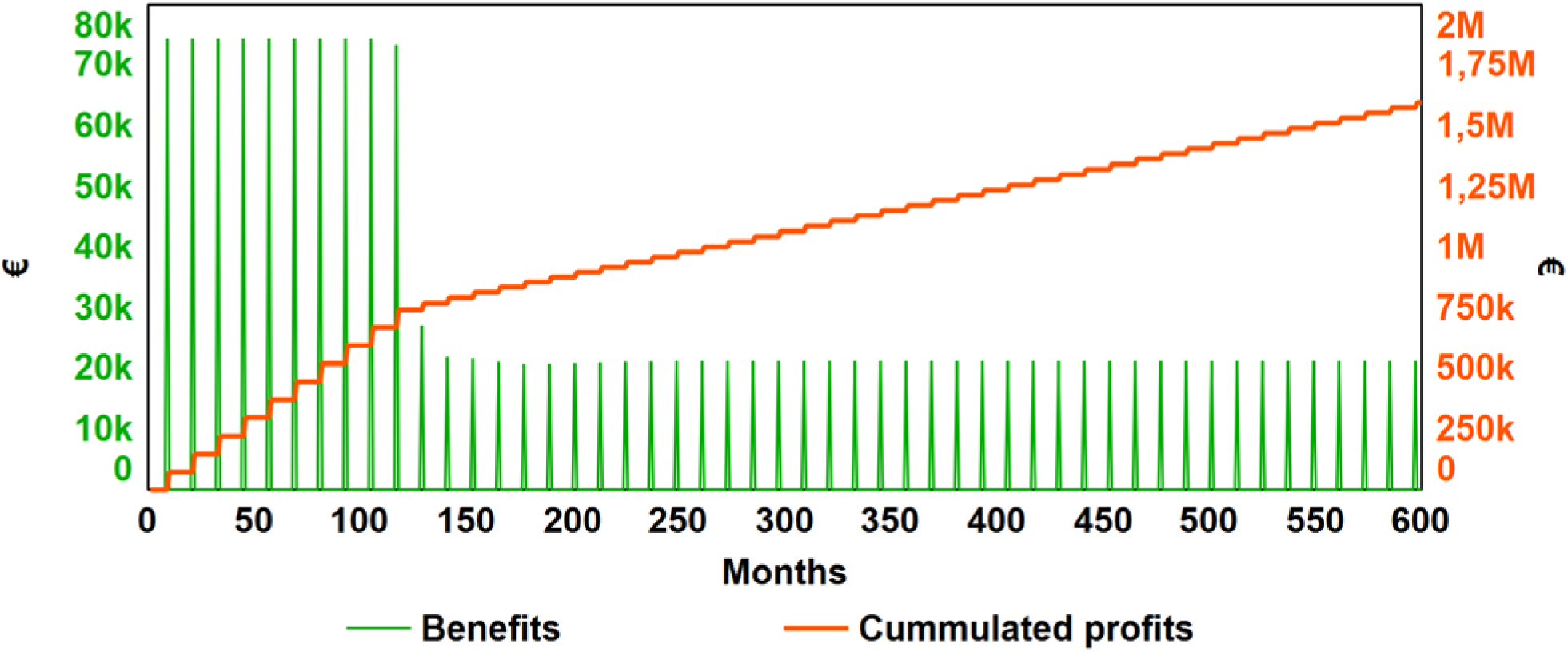
*Egeria densa* total biomass evolution during 50 years in the 1,290 hectare system.

## 4. *Egeria densa* removal strategies

One of the most employed strategies when dealing with invasive species, both in terrestrial and aquatic environments, is their removal (Bonanno, 2016; García-Díaz et al., 2020; Eppinga et al., 2020). In the case of *E. densa*, it is in part apparently very simple to carry out, since the floating portion of the plant can be eliminated with the use of nets or hooks. However, it is important to consider that there is a significant proportion of the plant that is attached to the substrate that can not be eliminated easily.

In this section, four strategies are compared in cost and efficiency in preserving the native ecosystem against the threat of the invasion. All the proposed strategies started at month 120 (year 10), when the density of *E. densa* reached 18% of the carrying capacity (Figure 10) and economic losses dropped by 64% in the following months. We established a variable removal price using a graphical function, with a higher price per kg extracted when low densities occurred (6 € kg^-1^) and a lower price per kg in high densities (0.3 € kg^-1^).

### Strategy 1: Yearly removal of floating E. densa during the peak abundance month

We simulated what would happen if different amounts of floating *E. densa* were removed from the system during October, when the abundance and density was higher during the year. We introduced parameters on the *Floating Removal Rate* variable to simulate a yearly removal of the 25%, 50% and 100% (Table 5). The changes observed (Figure 11) were not particularly significant in comparison to the effort and resources invested. After 50 years of simulation, the amount of *E. densa* was reduced by approximately by 21%, only achieved when 100% of the floating *Egeria* was removed. The accumulated costs associated to these strategies were 3 (-25%), 7 (-50%) and 24 million € (-100%), removing 6 (-25%), 7 (-50%) and 22 million Kg (-100%) after 40 years of actions.

**Figure 11.**
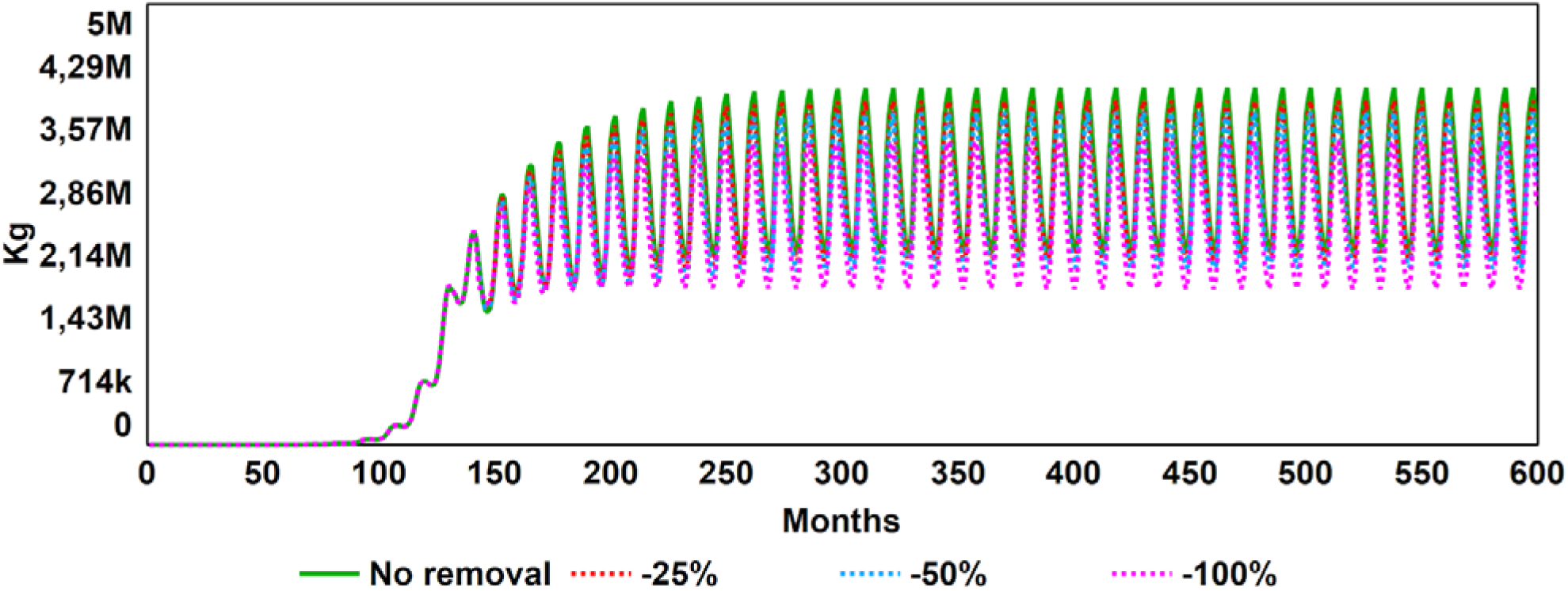
Comparison of removal strategies. Yearly removal of *floating Egeria densa* during each October, when the peak month occurs.

**Table 5.** *E. densa* removal strategies tested on the dynamic model.

| Parameter |  | Total removal rate | Floating removal rate |
| --- | --- | --- | --- |
| Base |  | 0 | 0 |
| Floating *E. densa* on the highest abundance month | -25% | 0 | x0.25 (October) |
|  | -50% | 0 | x0.5 (October) |
|  | -100% | 0 | x1 (October) |
| Total *E. densa* on the highest abundance month | -25% | x0.25 (October) | 0 |
|  | -50% | x0.5 (October) | 0 |
| Floating *E. densa* monthly removal | -10% | 0 | x0.1 (every month) |
|  | -25% | 0 | x0.25 (every month) |
|  | -50% | 0 | x0.5 (every month) |
| Total *E. densa* monthly removal | -5% | x0.05 (every month) | 0 |
|  | -10% | x0.1 (every month) | 0 |
|  | -20% | x0.2 (every month) | 0 |

### Strategy 2: Yearly removal of all Egeria densa during the abundance peak month

We tested what would happen if different amounts of both floating and planted *E. densa* were removed from the system, again during October, the month of maximum abundance. We introduced feasible parameters on the *Total Removal Rate* variable to simulate a yearly removal of the 25% and 50% (Table 5). The changes observed (Figure 12) were a reduction of the 35% of *E. densa* biomass when the 25% was removed, and a reduction of 58% when the 50% was removed. The accumulated costs associated with these strategies were 44 (-25%) and 119 million € (-50%), removing 31 (-25%) and 47 million Kg (-50%) after 40 years of actions.

**Figure 12.**
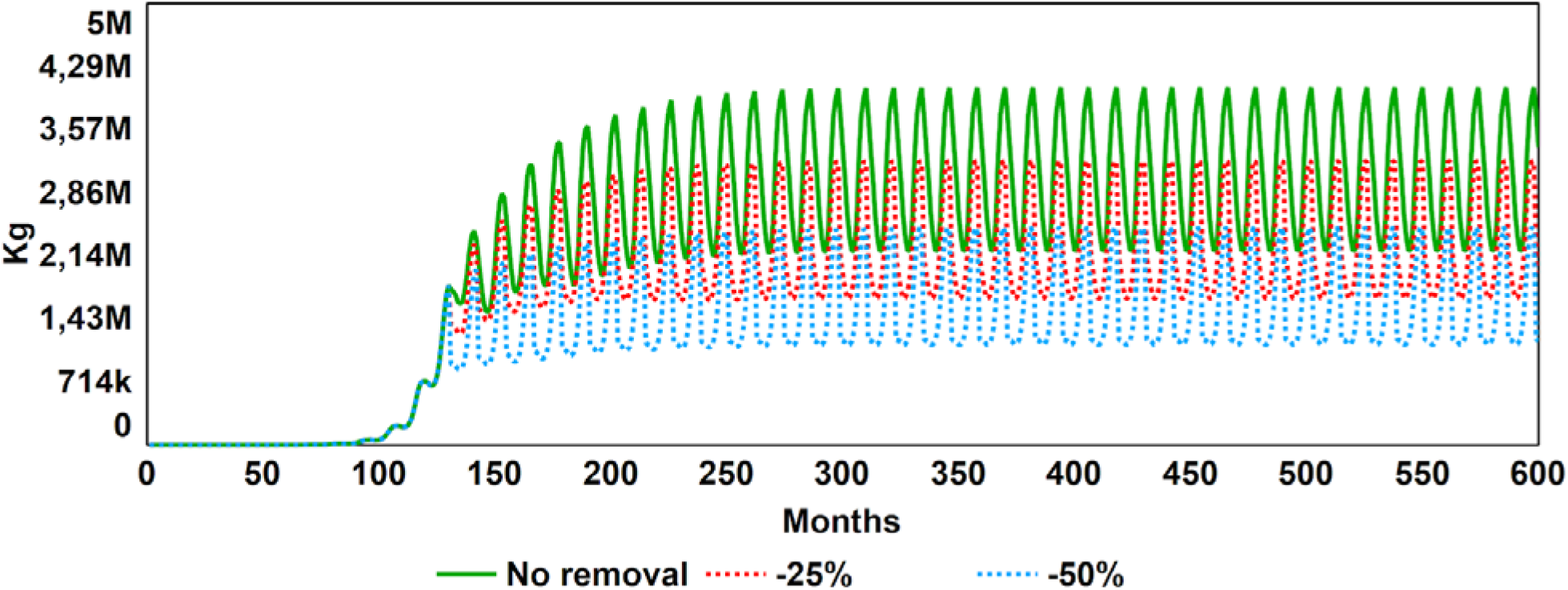
Strategy output from each percentual removal. Yearly removal of *Floating and Planted Egeria densa* during the peak month of abundance (October).

### Strategy 3: Monthly removal of floating Egeria densa

This strategy simulates what would happen if different amounts of *E. densa* were removed every month. We introduced parameters on the *Floating Removal Rate* variable to simulate a monthly removal of the 10, 25 and 50% (Table 5). The outputs showed (Figure 13) a reduction of the 15% (-10%), 27% (-25%) and 42% (-50%) of biomass when *E. densa* was removed. The accumulated costs associated with these strategies were 29 (-10%), 56 (-25%) and 107 million € (-50%), removing and 16 (-10%), 27 (-25%) and 40 million Kg (-50%) after 40 years of actions.

**Figure 13.**
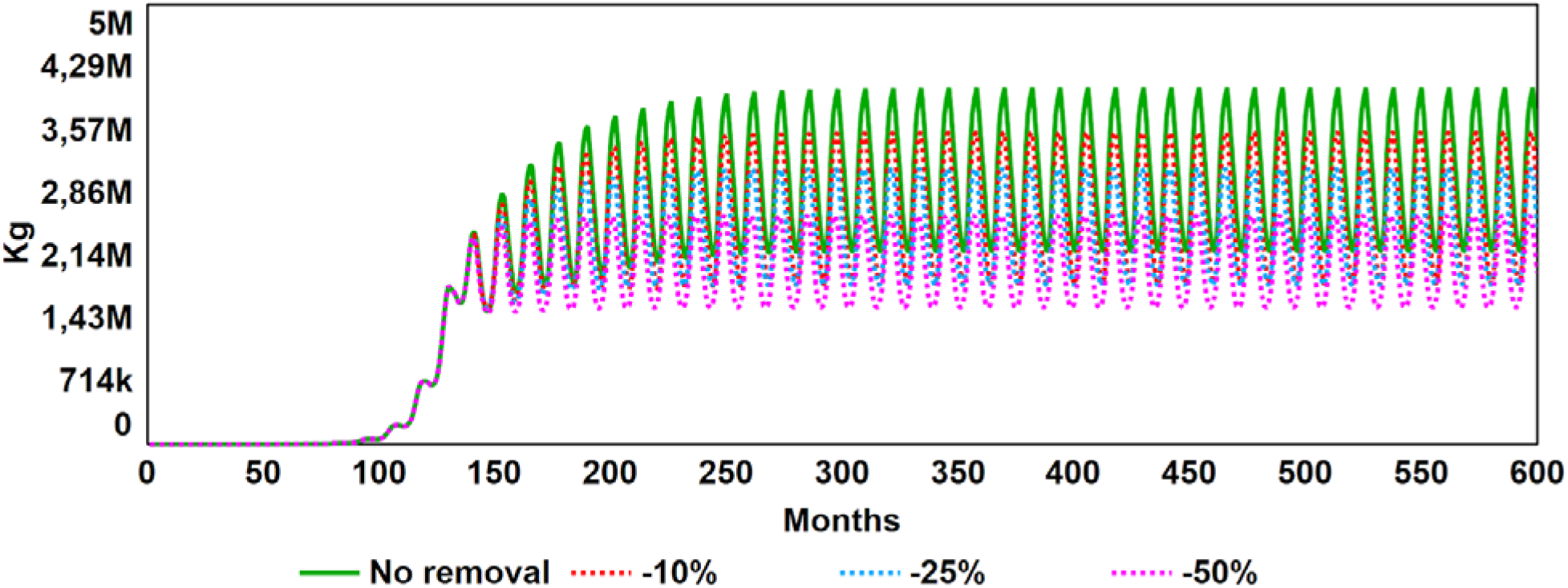
Monthly removal of *floating Egeria* strategy output from each percentual removal.

### Strategy 4: Monthly removal of all Egeria densa

This last removal strategy simulated what would happen if different amounts of *E. densa* were removed every month. We introduced parameters on the *Floating Removal Rate* variable to simulate a monthly removal of the 5, 10 and 20% (Table 5). The outputs showed (Figure 14) a reduction of the 53 (-5%), 84 (-10%) and 100% (-20%) when *E. densa* was removed. The accumulated costs associated to these strategies were 137 (-5%), 138 (-10%) and 8.83 million € (-20%), removing 43 (-5%) and 27 million Kg (-10%) after 40 years of actions, and 1.16 million Kg (-20%) after 5-6 years.

**Figure 14.**
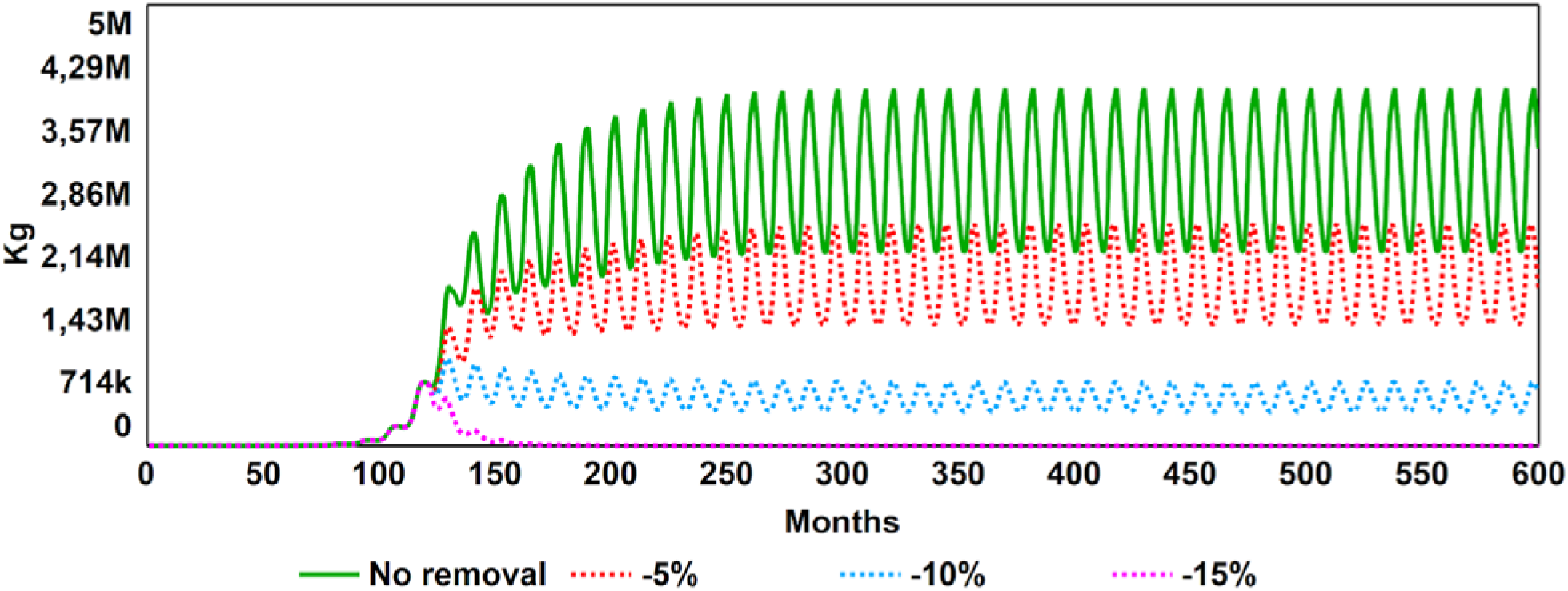
Monthly removal of *Floating* and *Planted Egeria densa* strategy output from each percentage of removal.

## 5. Discussion

The model output was able to reproduce the general dynamics of an invasion, showing a S-shape growing pattern. The addition of the carrying capacity submodule allowed it to mimic how the population continued its expansion until the intraspecific competition for resources, such as nutrients, light or space, stopped the exponential growth and reached a plateau (carrying capacity) (as seen in blue in Figure 9). Although we didn’t change the carrying capacity value (*k*), other scenarios could be built by modifying its value. The dynamics shown by the model were theoretical, since there are no historical time-series of the invasion process, nevertheless since the scenarios comparison are less sensitive to the growth rate, the results of what scenarios are better, would not change. Despite of this absence of field data, we managed to obtain a dynamic model that could be easily fitted to real data by changing the growth rates and the *k* and is similar to that of other aquatic invasive plants like the Japanese knotweed (*Fallopia japonica*) in Western Europe (van Oorschot et al., 2017).

The dynamic model comprises the entire *E. densa* life cycle, considering when the plant is attached to the bottom and when it is free living. The model should be considered as a frame to understand the dynamics of the invasion process of an aquatic plant and a tool to compare different management strategies, rather than replicating a concrete invasion process. For this reason, as far as the model reproduces a feasible output and responds to change of variables as expected, we consider the model a useful tool for managers even if it is not an accurate depiction of reality, in any case, the parameters values of the model could be easily changed to fit a concrete invasion process. The starting amount of *E. densa* (3 kg) was chosen emulating the release of the plants of an aquarium, it must said that the results will not differ from the simulations done if the amount of the initial biomass was half or double, due to the exponential growth of the species during the first years (10 years, Figure 9). The model only takes into account the asexual reproduction, although the sexual is designed but inactive, this is because the plants in the studied sites are all the same gender making sexual reproduction impossible, pointing out that the invasion could have happened by only one release and the existing individuals are clones.

However, we added in the model the possibility of adding a sexual reproduction, considering flowers stock with its seasonality and flowering rate. This part has not been used but can be activated by adding the parameter values.

The environmental variables in the model (temperature, light, nitrogen, and water turbidity) are those that affect *E. densa* growth and development. Other variables such as oxygen, pH, salinity, phosphate, or the amount of other nitrogen species could be added if the relationship with the growth rates were known, an aspect that could be explored in future research.

Regarding the economic output of the model, it was useful to simulate the effect on profits of the rice crops that the invasion had and compare different scenarios and the calculation of the costs of the removal strategies.

Removal strategies have been widely used in the management of aquatic exotic plants, such as *Ludwigia grandiflora* (Newman & Duenas, 2010; Hussner et al., 2016), *Hydrocotyle ranunculoides* (Kelly, 2006; Delbart et al., 2013) or *Crassula helmsii* (Delbart et al., 2013). Considering that sometimes these removals are not successful at all (Beric & Maclsaac, 2015) and they do not solve the problem, e.g. *Hydrilla verticillata* in Florida (WANS, 2003) or in California (Akers, 2012). According to our model’s output, the best strategy to remove *E. densa* would be a monthly removal of around 20% the total amount of the plant once it has been detected on the ecosystem (Figure 19). This management would lead to stopping the expansion and avoiding worse damage to the ecosystem. The cost of this optimal strategy is 8.83 million €. Comparing the different simulations performed, it looks better and cheaper to remove *E. densa* rapidly than letting it continue with its expansion. That is because if it is not managed like this, the population would continue with its exponential growth and any of the proposed management would not be effective to act in benefit of the health status of the ecosystem as it would be more expensive.

**Figure 19.**
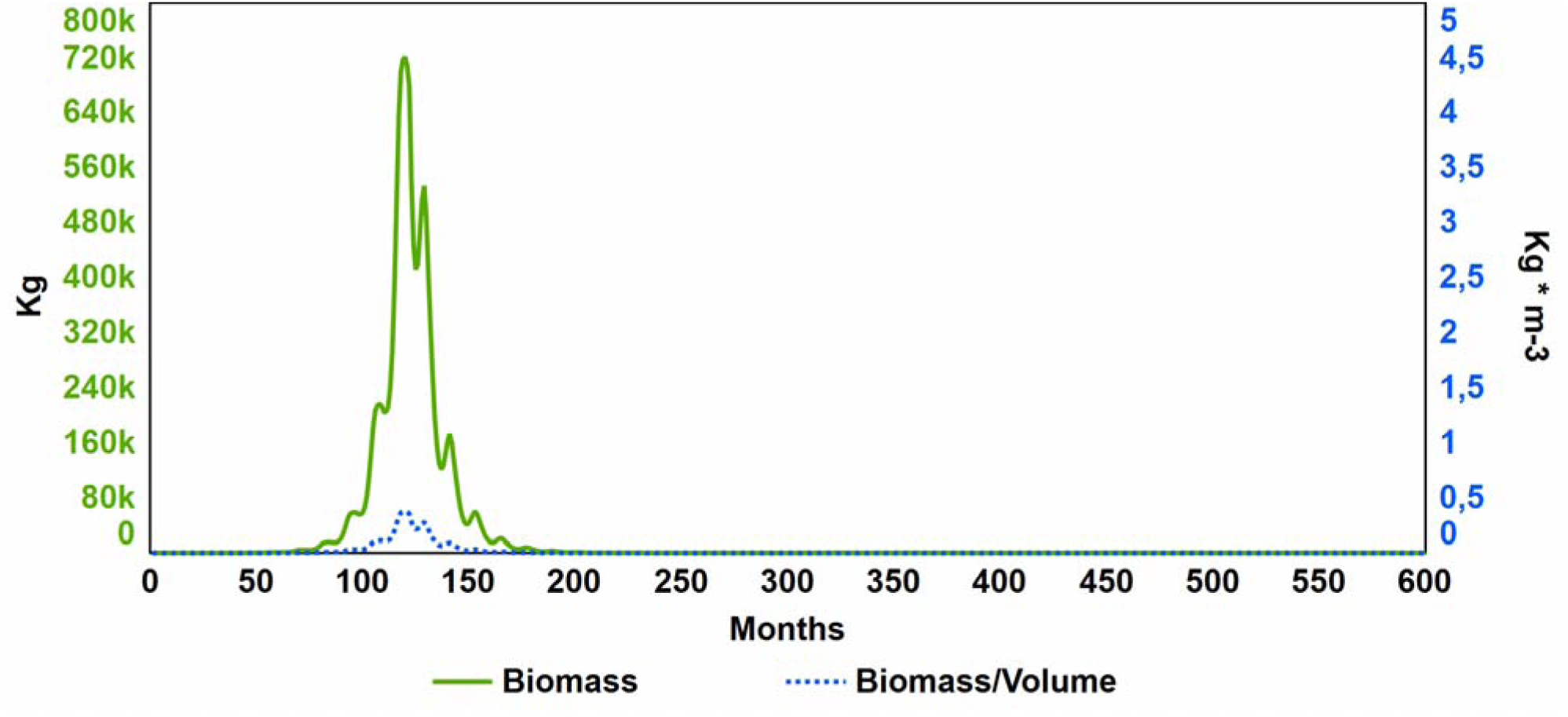
*Egeria densa* total biomass evolution during 50 years in the 1,290 hectare ecosystem when the 20% monthly removal of the plant is performed (part of scenario 4).

On the other hand, the removal of floating *Egeria* is not effective enough to compensate for the cost of the management done in that way. It will always be more effective the removal of planted and floating *E. densa* because the problem lies on the fragmentation of the one that is attached to the bottom which would continue with its constant release.

There is still an enormous variety of other possible combinations in removal strategies to explore, only a few of them are evaluated in this work, but the model done could be adapted to any scenario. Other interesting strategies used in management of aquatic invasive plant species are nutrient reduction (Wittmann et al., 2015; Coetzee & Hill, 2012); chemical control using herbicides (Getsinger et al., 2008) or salty water (Thouvenot et al., 2012); biologic control introducing insects (Cuda et al., 2008), fungal pathogens (Shearer, 2013) or herbivores such as the grass carp (Pipalova, 2006); or other more ingenious ones as the dye application to avoiding light reaching photosynthetic pigments (Bornette & Puijalon, 2011); water level drawdown in channels during maintenance periods (Dutartre et al., 2006); and other strategies reviewed in Hussner et al., 2017.

Finally, the use of population models for different aspects such as determining growth or expansion, or for advising on invasive species decisions, is not new. An example would be the models developed for Japanese Knotweed (*Fallopia japonica*) and Salicaceae species where vegetation colonisation, growth, prediction of hydraulic resistance, and mortality through flooding, desiccation, etc. are monitored (van Oorschot et al., 2017). In the specific case of *E. densa*, there are a variety of models that explore different aspects of the ecology of the species: temperature, water turbidity and the presence of surfactants as growth factors (Machado et al., 2020; Wanderley et al., 2021); Cu and Zn ions bioremediation capacity (Módenes et al., 2012); the regulation of its photosynthesis (Laing & Browse, 1985); or its biomass distribution, growth and decay (Carrillo et al., 2006). The contributions of our model start from the imitation of the dynamics and the inclusion of environmental variables, which had already been modelled so far, being the most outstanding novelties, the simulation of removal strategies and their effect on profits of rice crops and cost of removal.

## 6. Conclusion

This research develops a dynamic model which recreates the invasion of an aquatic alien plant. According to the results, the optimal strategy for maintaining the ecosystem in the best condition possible is to eliminate the *E. densa* floating shots as soon as possible, avoiding the high growth speed reached after some years after the first colonisation.

